# Sea urchins, parrotfish and coral reefs in Grand Cayman, BWI: exemplar or outlier?

**DOI:** 10.1101/2020.12.11.421867

**Authors:** Elizabeth Sherman

**Affiliations:** Natural Sciences, Bennington College, Bennington, Vermont, 05201, USA

## Abstract

The change in state of Caribbean coral reefs over the last 40 years has been characterized by phase shifts from scleractinian coral cover to macroalgal cover, the loss of structural complexity and a decline in biodiversity. Not only do scientists want to understand these changes, but also predict the future of coral reefs and their capacity for resilience. In particular, the loss of herbivory, due to declines in parrotfish and the sea urchin *Diadema antillarum*, has been implicated in many studies as a proximate cause of the coral to macroalgal phase shift. However, reports of the particular role of these putative herbivores have varied, with some studies claiming a causal role for parrotfish, others for *Diadema* and still others suggesting no such relationships. Often these studies just examined one response measure of coral reef biodiversity. In this paper, I report the relationship between parrotfish and *Diadema* to many metrics of reef organization surveyed simultaneously in the same transects in reefs outside and within the Marine Protected Area (MPA) of Grand Cayman, an island that has been affected by increasing tourism over the last 30 years. The magnitudes of the various measures of reef diversity reported here are consistent with those reported elsewhere. The relationships among those measures are consistent with those reported in some prior studies and inconsistent with others, reflecting the variation in responses documented in prior studies. The presence of sea urchins was associated with survey sites having higher levels of coral cover, lower levels of macroalgae cover, and lower densities of parrotfish than survey sites without sea urchins. Moreover, parrotfish abundance was associated with a decrease in coral cover and little relationship to macroalgae cover. Neither coral cover nor macroalgae cover was different in sites within the MPA compared to sites outside the MPA. I argue that the combination of site-specific local stressors and their interaction with global stressors makes it unlikely that any one island or even regional reef system could serve as an exemplar for Caribbean-wide reef degradation. Moreover, it is difficult to assess the potential for reef resilience in the face of the ongoing assaults from increasing tourism pressures and global climate change.

## INTRODUCTION

Over the last forty years, coral reefs of the Caribbean have experienced widespread phase shifts from stony coral dominance to macroalgal dominance with a concomitant decrease in reef structure, function and biodiversity (Adam et al, 2015; Bruno, Coté, and Toth, 2019; Shantz, Ladd and Burkpile, 2020). It is well-established that macroalgae out-compete coral for substrate and light, through a variety of mechanisms including their more rapid growth rate, mechanical and allelochemical inhibition, and alteration of the benthic microbiome (Arnold, Steneck, and Mumby, 2010; Box and Mumby, 2007; Foster, Box, and Mumby, 2008; Thurber et al., 2012). The proximate causes that have been linked to the coral-to-macroalgal phase shift include the loss to overfishing or disease of the herbivores that would otherwise graze on macroalgae. These proximate causes are likely exacerbated by additional stressors, both local and global (Adam et al., 2015; Graham et al., 2015; Mora, 2008). The two main groups of animals associated with coral reef herbivory in the Caribbean are sea urchins, in particular the long-spined sea urchin, *Diadema antillarum*, and parrotfish. In the mid-1980s, a still-unidentified disease caused a massive Caribbean-wide die off of *D. antillarum*, which prior to that, had been implicated in significant macroalgal grazing (Lessios, 2016). Following that die-off, phase shifts from coral-dominated to macroalgae-dominated reefs were widely reported (Lessios, 2016). Additionally, parrotfish have been reported to graze on macroalgae and their loss due to overfishing has been associated with macroalgal proliferation (Mumby, 2009; Mumby et al, 2006). Nevertheless, in spite of ongoing investigation, questions regarding the specific functional roles of these two groups of herbivores persist (Adam et al., 2015). Early studies reported correlations between parrotfish density, parrotfish grazing intensity and coral cover (Mumby, 2009; Mumby et al, 2006). Results from more recent experimental studies in which parrotfish of various sizes were excluded from settling plates in various parts of Caribbean reefs were consistent with a top-down causal relationship between parrotfish grazing and coral cover (Burkpile and Hay, 2010; Steneck, Arnold, and Mumby, 2014). Others, however, failed to find such a relationship and, in fact, suggested that it is the benthos that influences parrotfish density (Suchley, McField, and Alvarez-Filip, 2016). Furthermore, Clements et al. (2017) argued that parrotfish are microphages and play little role in consuming macroalgae. At the same time, other studies reported that increased densities of *D. antillarum* were associated with decreases in macroalgal density (Maciá, Robinson and Nalevanko, 2007; Myhre and Acevedo-Gutiérrez, 2007) and concomitant increases in juvenile coral abundance (Carpenter and Edmunds, 2006; Edmunds and Carpenter, 2001; Idjadi, Haring, and Precht, 2010). Nevertheless, other reports found no such relationship (Lacey, Fourqurean and Collada-Vides, 2013; Mercado-Molina et al., 2015; Rodríguez-Barreras et al, 2018; Steiner and Williams, 2006).

More recent studies have endeavored to tease out the variables that affect herbivory, macroalgae, and coral cover, and have revealed complex relationships among the functional roles of different herbivores, even within the same taxonomic group. Adam et al (2018) reported that various species of parrotfish functioned as scrapers, excavators, croppers or macroalgae browsers and Shantz, Ladd, and Burkepile (2020) further demonstrated that parrotfish of different sizes also partitioned the benthic resources in coral reefs.

In 2008, Knowlton and Jackson called for empirical studies to establish baselines of coral reef structure, function and biodiversity so that their changes could be monitored, knowing that those baselines were and would continue to be moving targets. Bellwood et al. (2019) reiterated that charge arguing that high resolution data at different scales of reef ecosystem function (local, regional, global) are still lacking as is a more complete understanding of the extent to which herbivore functions are complementary or redundant (Lefcheck et al, 2019). These data are essential in order to understand the persistent uncertainties regarding coral reef resilience (Bruno, Coté, and Toth, 2019).

In this study, I recorded data of both explanatory variables and response variables of reef biodiversity simultaneously in the same reefs surrounding Grand Cayman, the largest and most populous island of the Cayman Islands, BWI. The value of this research is that various aspects of benthic cover were surveyed simultaneously with possible explanatory variables in different locations (within the Marine Protected Area or not) at different depths in the same transects on various reefs around Grand Cayman. Studies measuring both sea urchin density and parrotfish size distribution and density along with measures of benthic composition (coral cover, macroalgae cover, turf algae cover, crustose coralline algae [CCA] cover) simultaneously in the same sites are lacking and can facilitate untangling the functional roles of these animals in the coral reef ecosystem.

Like many islands in the Caribbean, Grand Cayman is a major tourist destination with growing infrastructure and concomitant assaults on the reef (Hurlston-McKenzie et al., 2011). The intensity of the tensions between supporters of environmental protection and those supporting massive shore and land development are particularly apparent and heated. For example, in 2013, the National Conservation Law was established to expand protection of critical habitats and organisms. Yet the ongoing development of the island seems inconsistent with that Law (e.g. Lederer, 2019; Whittaker, 2020). The Cayman Islands Department of Environment has pushed for the expansion of reef areas to be included in the MPA at the same time that the Legislative Assembly approved a proposal for a new cruise berth facility that would require dredging of George Town Harbor (Nunis, 2019), spurring the emergence of a grass roots movement opposing the facility. These tensions are only likely to increase in the future making the present study of reef diversity in Grand Cayman all the more critical. Can reef resilience be promoted under these conditions? Moreover, as reefs continue to be stressed and assaulted both locally and globally, what is the state to which we hope they return?

## MATERIALS AND METHODS

### Study Sites

I conducted surveys in the fringing reefs located in western and northern areas around Grand Cayman, Cayman Islands, B.W.I. (19.3222° N, 81.2409° W), as they were reliably accessible by boat and scuba. The leeward side of Grand Cayman is the western coast and the near-shore environment is distinguished by ironshore or sand, contiguous with the land, followed by the reef flat further from shore that is largely composed of sand and reef debris. That then leads to a shallow coral reef terrace (depths from roughly 5-12 m) characterized by spur and groove formations as well as more discrete coral patches. The shallow terrace leads to a sand flat lagoon and then to the deeper coral reef shelf at depths of about 16-25 m from which the shelf descends down into abyssal depths (McCoy, Dromard, and Turner, 2010). The northern reef areas are windward, with reef complexes similar to those on the west, except that the wall from the deeper reef descends more steeply. Moreover, the western reefs I surveyed were in the Marine Protected Area (MPA), while the northern sites were not (Fig. 1).

**Figure 1.**
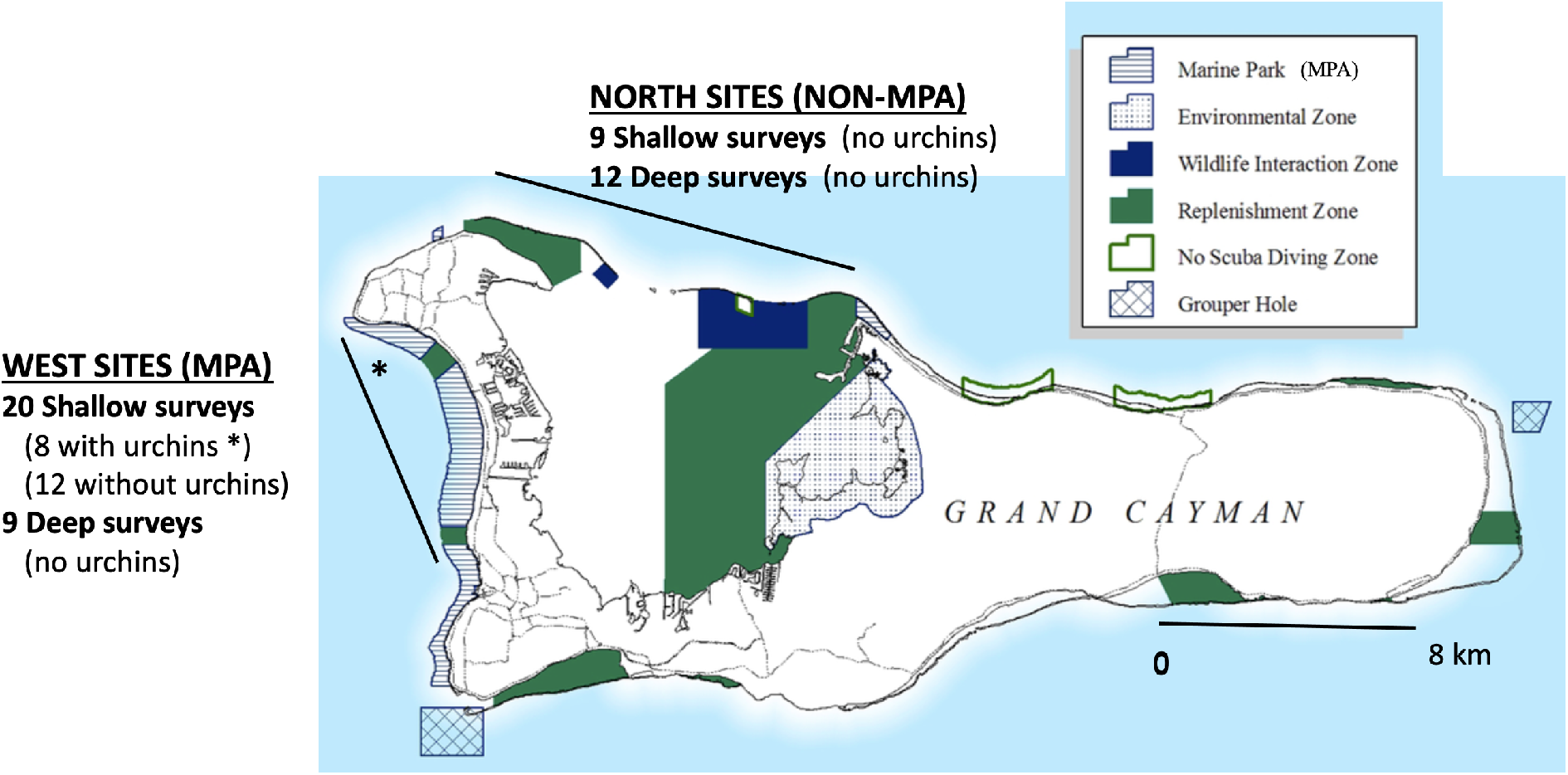
Map of marine park designations of Grand Cayman. The dark straight lines on the west and northwest represent the areas of the survey sites. The asterisk indicates the small sequestered area having sea urchins. See text for further explanation of the survey sites. http://doe.ky/marine/marine-parks/

*Diadema antillarum* (hereafter, *Diadema* or sea urchins) are distributed heterogeneously around the island. Large populations can be reliably found in small hardpan ironshore inlets right next to shore (Bellwood et al’s so-called sea urchin barrens, 2004; Fig. 2) but I have never found them in great numbers in actual living coral reef for the previous 15 years or so, although they were once abundant (at depths from 5 m to > 20 m) prior to the great die-off of the 1980s (Fig. 1S-supplementary material).

**Figure 2.**
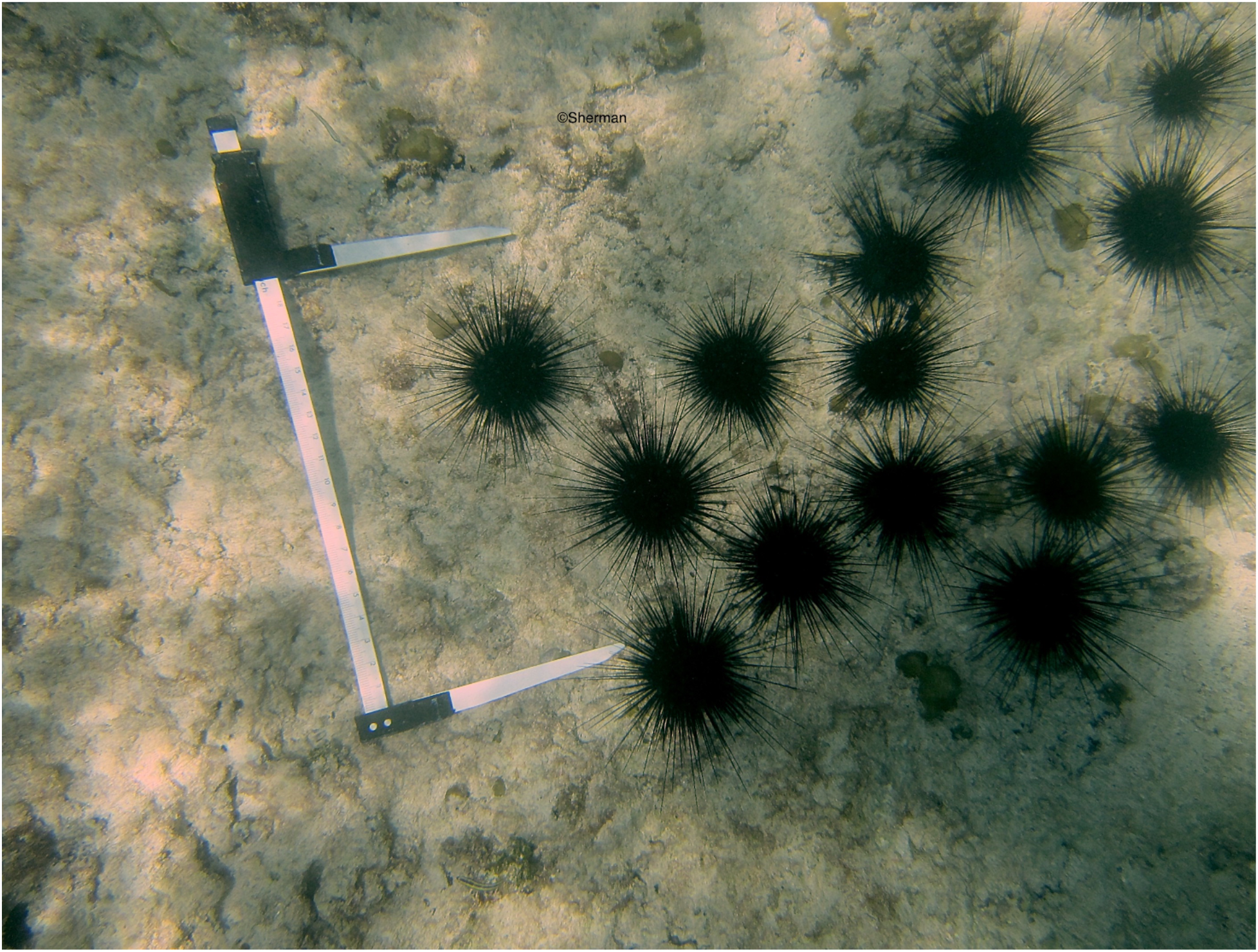
Typical density of *D. antillarum* in hardpan inlet (not reef). Note that much of the hardpan is bare.

A serendipitous finding of sea urchins inhabiting coral reef proper in a circumscribed part of the shallow western reef terrace within the MPA permitted a part of this study in which I compared benthic diversity and parrotfish diversity in reefs with and without sea urchins present. Typically, when one descends into the reef using scuba, the reef appears fuzzy, which is due to an abundance of macroalgae, predominantly *Dictyota sp.*, covering the reef (Fig. 3a). But the first time I dove in one of the survey sites that turned out to have sea urchins, the reef did not look fuzzy, but rather appeared to have discrete edges and boundaries (Fig. 3c). When I got closer, I saw that there was very little algae and noticeable numbers of sea urchin (compare Figs. 3b, close-up of site without sea urchins, and 3d, close-up of site with sea urchins; see videos in supplemental material: chain no urchins.mv4; chain with urchins.mv4). These survey sites have been the only reef sites in which I have found *Diadema* at measurable densities to date. They were roughly 15-30 m away from other parts of the reef that had no sea urchins. In some analyses, the surveys from those 8 sites are compared to the other 42 that lacked visible sea urchins. The reef survey sites with sea urchins were not scattered randomly throughout the western shallow reefs, but rather sequestered in one area. The area with the sea urchins (located in the more northwestern part of the western shallow reefs, Fig.1) is also smaller compared to the area without sea urchins.

**Figure 3.**
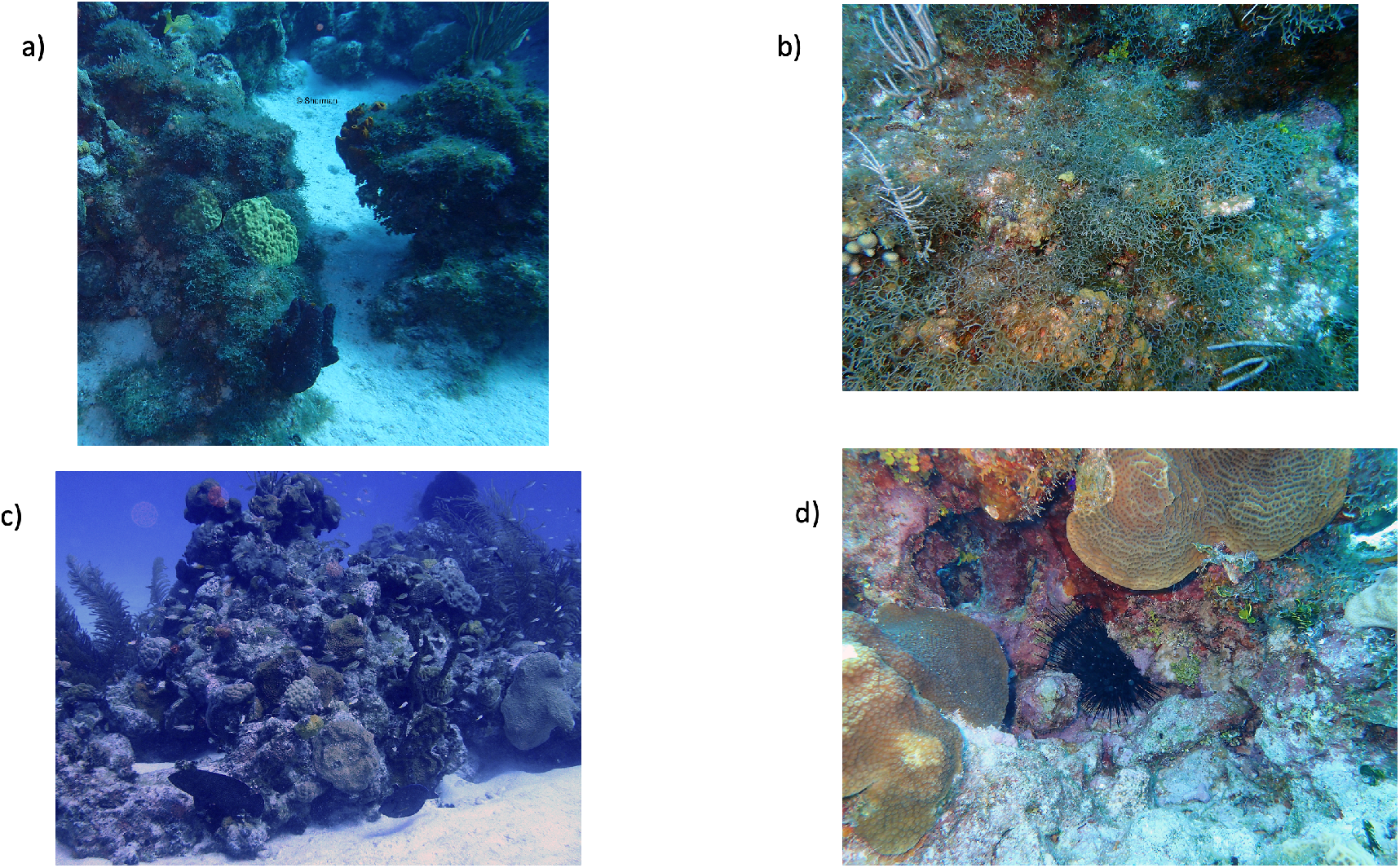
Exemplary photos of reefs in the West-shallow survey sites with and without sea urchins. a) Typical reef with dense macroalgae cover. b) Close-up of reef with dense macroalgae cover. c) Reef with low macroalgae cover in an area with *Diadema*. d) Close-up of reef with low macroalgae cover in an area with *Diadema*.

In summary, 50 surveys were conducted distributed across the various locations. There were 28 individual survey sites from the West, which is part of the MPA, and leeward: 19 shallow survey sites (depths of 9-14 m), 8 of those survey sites with urchins present, 11 without, and 9 deep survey sites (depths of 18-24 m), none of which had sea urchins. Twenty-two surveys were conducted in individual sites from the North, which is windward and not within the MPA: 9 shallow survey sites (depths of 9-14 m) and 13 deep survey sites (depths of 18-24 m), none of which appeared to have sea urchins (Fig. 1). The surveys were conducted over a period of 18 months in July 2017, July 2018, and January 2019.

### Surveys

At each location of the 50 survey sites, underwater transect plots were chosen haphazardly. After descending to the reef using scuba, I swam along the reef and spooled out a 30 m transect line while conducting underwater visual censuses of parrotfish (in an area 2 m on either side of the transect line), estimating total parrotfish length in ranges of <11 cm, 11-20 cm, 21-30 cm, 31-40 cm, >40 cm. Prior to the censuses, I had done underwater trial sessions with objects of known lengths in order to correct for the underwater length visual distortion. The five most abundant parrotfish at all survey sites were the stoplight parrotfish (*Sparisoma viride*), redband parrotfish (*Sp. aurofrenatum*), striped parrotfish (*Scarus iseri*), princess parrotfish (*Sc. taeniopterus*), and queen parrotfish (*Sc. vetula*). Very occasionally itinerant yellowtail (*Sp. rubripinne*), redtail (*Sp. chrysopterum*), and rainbow parrotfish (*Sc. guacamaia*) would traverse the transect and were noted but due to their unreliable and infrequent siting, they were not included in the quantitative analyses. Following the parrotfish census, I placed a 25 cm x 25 cm quadrat, every 5 m along the 30 m transect line, and photographed the quadrat for a record of the benthos (1 m above the quadrat), for later analysis. Finally, a research assistant and I counted the number of *D. antillarum* within 5 m on either side of the 30 m transect. Thus, I had surveys of parrotfish, benthic cover, and sea urchins within each 30 m transect.

### Analysis of benthic photographs

There were 6 quadrat photos for each of the 50 transects that were surveyed. Each photo was opened in ImageJ (Abràmoff, Magalhães, and Ram, 2004), and a 6 x 6 grid was superimposed (Fig. S2). The benthos under each of the 36 points of intersection was identified as living stony coral, fleshy macroalgae, turf algae, CCA, or other categories not reported here (including octocorals, sponges and benthos that could not be identified, the latter comprising less than 1% of the points). Individual species within the categories were not recorded. For each of the 50 transects of the survey sites, I computed the average per cent cover of the various benthic categories for the 216 points.

### Data analysis

In order to assess the effect of survey sites on the different response measures of reef diversity, I performed two-way ANOVAs with survey site category and year as predictor variables. In addition, I wanted to determine if the sea urchin survey site data should be analyzed as a separate category, so initial two-way ANOVAs were performed without data from sea urchin survey sites, and then again with sea urchin survey site data added to the rest of the shallow western survey site data (supplementary material: Two-way ANOVAs). If survey site category was not significant without the sea urchin survey site data but became significant when sea urchin survey data were added, I treated the sea urchin survey sites as a separate site category in subsequent analyses. Additionally, “year” was not a significant predictor of most of the response variables in the two-way ANOVAs, so I pooled data from the three years in subsequent one-way ANOVAs. These ANOVAs tested whether the five different categories of survey sites (West-shallow with urchins, West-shallow without urchins, West-deep without urchins, North-shallow without urchins, North-deep without urchins) were associated with different magnitudes of response variables.

In order to test if sea urchin density was correlated with different measures of reef biodiversity (e.g. coral cover, macroalgae cover, CCA cover, turf algae, parrotfish abundance), I performed linear regression analyses. Because sea urchins were found only in a circumscribed area of shallow reefs on the West, I analyzed sea urchin density as a predictor of biodiversity measures from the two categories of Western-shallow reef survey sites only (with and without sea urchins). The rest of the regression analyses not involving sea urchin density as a predictor *per se* were performed using all the survey data from all the sites.

Finally, I examined parrotfish size distribution for each of the different categories of survey sites (separating the Western-shallow sites into those with and without sea urchins) using the pooled data for the 18 months of study.

## RESULTS

### Do measures of reef diversity vary with survey site?

Survey sites with sea urchins were associated with significantly higher percent coral cover than sites without sea urchins (Fig. 4a). A Tukey’s *post hoc* multiple comparison test revealed that the only significant comparisons were between the West-shallow sites with urchins and each of the other survey site categories, but there were no significant differences among the four other survey site categories. Moreover, survey sites with sea urchins were associated with significantly lower levels of macroalgae cover (Fig. 4b). Again, a subsequent Tukey’s multiple comparison test revealed that the only significant comparisons were between the West-shallow sites with urchins and each of the other site categories, but there were no significant differences in macroalgae cover among the four other survey site categories. In addition, while parrotfish density is potentially a predictor of coral reef cover and macroalgal cover, I tested it as a response variable and a one-way ANOVA revealed that survey sites with sea urchins exhibited a lower density of parrotfish than sites without sea urchins (Fig. 4c). The *post hoc* Tukey’s multiple comparison test demonstrated that, as with coral cover and macroalgae cover, the only significant comparisons were between the West-shallow sites with urchins and each of the other site categories, but there were no differences in parrotfish density among the four survey sites without sea urchins. Thus, survey sites with sea urchins were associated with higher levels of coral cover, lower levels of macroalgae cover, and lower parrotfish density compared to all the other survey sites (Table 1).

**Table 1.**
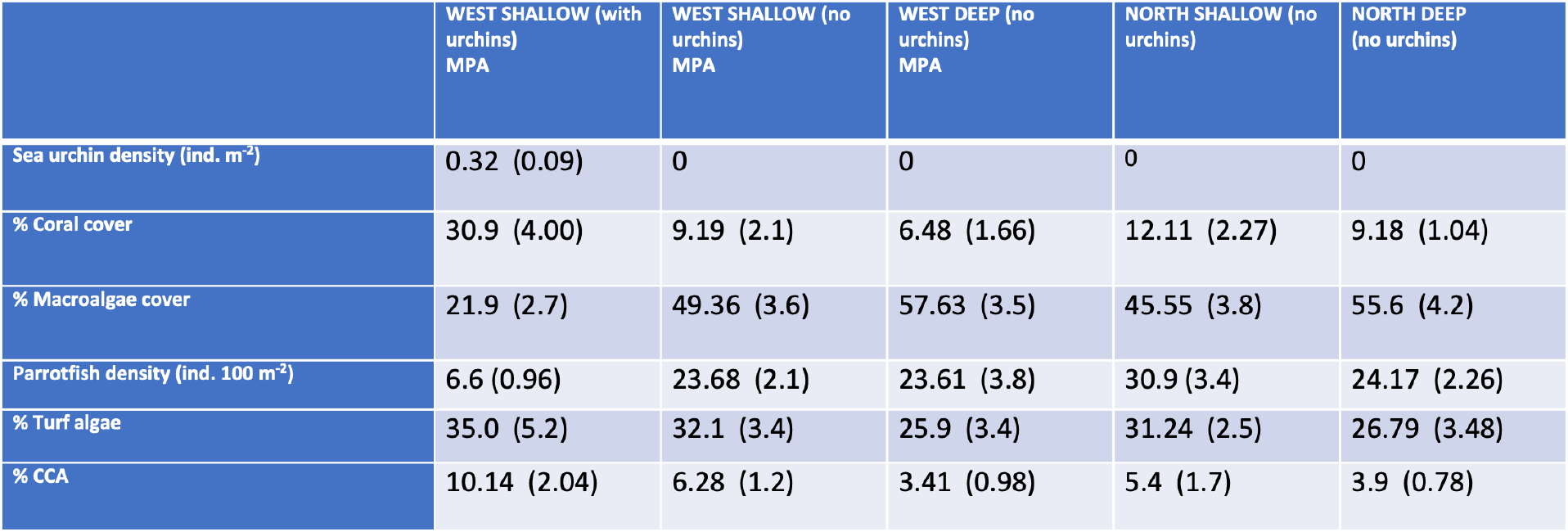
Mean coral reef diversity measures (SE in parentheses) in the 5 different categories of survey sites.

|  | WEST SHALLOW (with urchins)<br>MPA | WEST SHALLOW (no urchins)<br>MPA | WEST DEEP (no urchins)<br>MPA | NORTH SHALLOW (no urchins) | NORTH DEEP (no urchins) |
| --- | --- | --- | --- | --- | --- |
| Sea urchin density (ind. m<sup>-2</sup>) | 0.32 (0.09) | 0 | 0 | 0 | 0 |
| % Coral cover | 30.9 (4.00) | 9.19 (2.1) | 6.48 (1.66) | 12.11 (2.27) | 9.18 (1.04) |
| % Macroalgae cover | 21.9 (2.7) | 49.36 (3.6) | 57.63 (3.5) | 45.55 (3.8) | 55.6 (4.2) |
| Parrotfish density (ind. 100 m<sup>-2</sup>) | 6.6 (0.96) | 23.68 (2.1) | 23.61 (3.8) | 30.9 (3.4) | 24.17 (2.26) |
| % Turf algae | 35.0 (5.2) | 32.1 (3.4) | 25.9 (3.4) | 31.24 (2.5) | 26.79 (3.48) |
| % CCA | 10.14 (2.04) | 6.28 (1.2) | 3.41 (0.98) | 5.4 (1.7) | 3.9 (0.78) |

**Figure 4.**
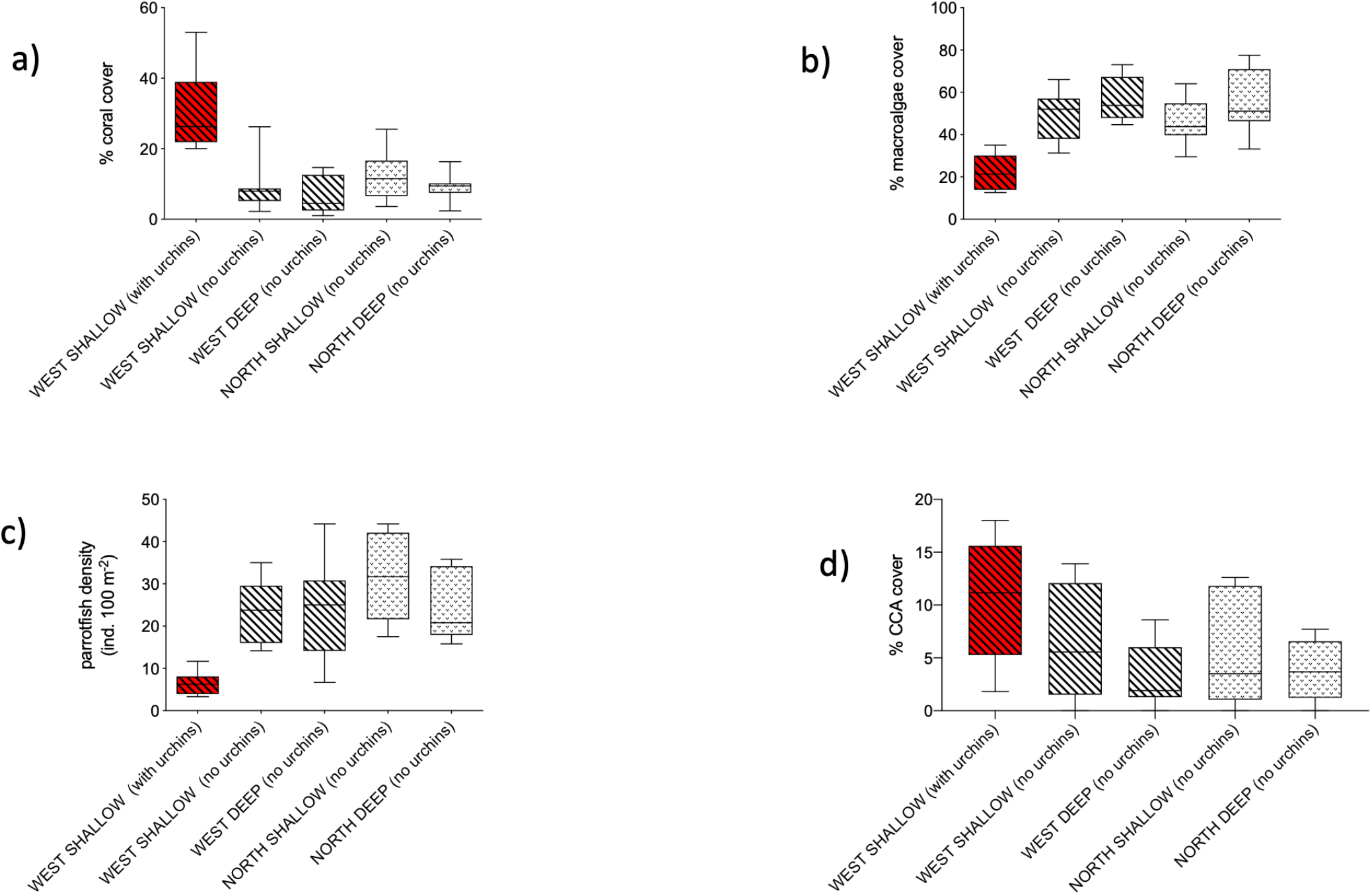
Different metrics of coral reef diversity as a function of the 5 different categories of survey sites: West-shallow with urchins (in red), West-shallow no urchins, West-deep no urchins, North-shallow no urchins, North-deep no urchins. The percent coral cover a), percent macroalgae cover b), and parrotfish density c) of the West-shallow survey sites with sea urchins were significantly different from those of the other 4 site categories without sea urchins (p<0.0001). *Post hoc* Tukey’s multiple comparison tests revealed that the only significant comparisons were between the West-shallow survey sites with urchins and each of the other survey site categories, but there were no significant differences among the four other survey site categories. The percent CCA cover d) of the West-shallow survey sites with sea urchins was significantly greater than that of the other 4 site categories without sea urchins (p=0.0132). However, the result of the *Post hoc* Tukey’s multiple comparison test demonstrated that the only significant comparisons were between West-shallow survey sites with urchins and West-deep survey sites and North-deep survey sites, both without urchins.

Percent CCA cover was significantly greater in survey sites having sea urchins compared to the other sites (Fig. 4d, Table 1). However, the result of the *post hoc* Tukey’s multiple comparison test demonstrated that the only significant comparisons were between West-shallow survey sites with urchins and West-deep survey sites and North-deep survey sites, both without urchins. The results of the original two-way ANOVAs for turf algae cover did not permit separating the sea urchins survey sites into their own category (supplementary material: Two-way ANOVA Tables 5Sa and 5Sb). Site category was not a significant predictor of turf algae cover with or without the addition of the sea urchin survey data. Moreover, turf algae cover varied with year. Nevertheless, I included mean percent turf algae cover across the 5 different categories of survey sites in Table 1 for comparisons to the other response variables.

The western survey sites were within the MPA while the northern sites were not. The comparisons among the four sites without sea urchins revealed that the location in or out of the MPA was not related to any differences in reef diversity measures (Table 1).

### Correlations between various measures of coral reef diversity

Given that the presence of sea urchins was associated with higher percent coral cover, lower percent macroalgae cover, and lower parrotfish density, I tested if there was a quantitative relationship between density of sea urchins and each of those response variables. The only areas with sea urchins present were in a circumscribed portion of the West-shallow survey sites, and thus I could not compare survey sites from all locations (West-shallow, West-deep, North-shallow, North-deep) with and without sea urchins. However, regressing percent coral cover on sea urchin density, comparing data only from combined shallow western survey sites (those with and those without sea urchins), revealed that sea urchin density was positively correlated with percent coral cover (Fig. 5a, p<0.0001). I repeated the regression analysis using only survey data from the sea urchin sites in case the sea urchin site category was a unique category separate from the other West-shallow sites and the relationship remained significant (p=0.043). Sea urchin density was negatively correlated to percent macroalgal cover using data from combined Western-shallow survey sites (p=0.02, Fig. 5b) but data from sea urchin sites alone were not predictors of macroalgal cover (p=0.33). That is, the correlation disappeared when all the points with a sea urchin density = 0 were eliminated. Figure 5c shows the relationship between sea urchin density and parrotfish density and includes data from all the West-shallow survey sites.

**Figure 5.**
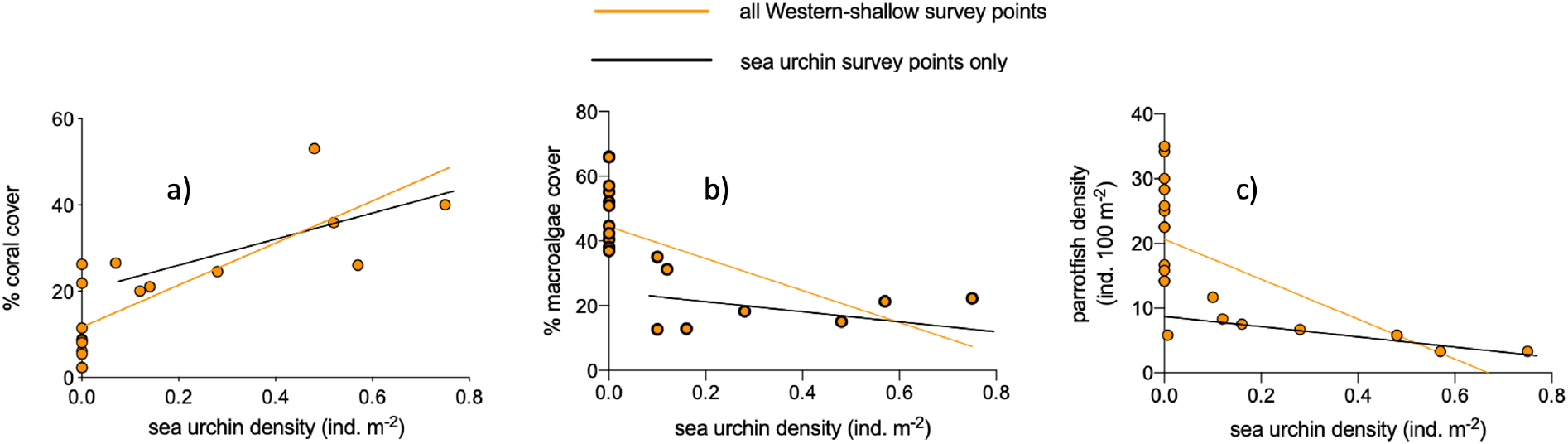
Sea urchin density as a predictor of three coral reef diversity measures using either all points from both West-shallow survey sites, with and without sea urchins (orange regression lines) or using data only from the sea urchin survey sites (i.e. not including the West-shallow sites with sea urchin density = 0, black regression lines). a) Using all points, the relationship between sea urchin density and percent coral cover was significant (orange line; p<0.0001); using data only from the sea urchin survey sites, the relationship remained significant (black line; p=0.043). b) Using all points, the relationship between sea urchin density and percent macroalgae cover was significant (orange line; p=0.003); using data only from the sea urchin survey sites, the relationship was no longer significant (black line; p=0.33). c) Using all points, the relationship between sea urchin density and parrotfish density was significant (orange line; p=0.0016); using data only from the sea urchin survey sites, the relationship remained significant (black line; p=0.036).

The relationship was significant whether the regression was performed with sea urchin survey data combined with the rest of the West-shallow survey site data (p=0.0016) or with the sea urchin survey site data only (p=0.036). Thus, sea urchin density was inversely related to parrotfish density. While percent CCA cover was significantly greater in survey sites having sea urchins compared to the other sites (Fig. 4d) there was no correlation between sea urchin density and percent CCA cover (Table 2). Finally, sea urchin density was not a predictor of turf algal cover (Table 2) if site data from both West-shallow survey data with and without sea urchins were used (p=0.36), and not quite significant when the West-shallow sea urchin survey data were tested alone (p=0.0504). The importance of this result is not straightforward as there was no significant difference in turf algae cover among the different site categories in the Two-way ANOVA (supplementary material: Table 5Sb).

**Table 2.** Relationships between the various measures of biodiversity (relevant figures or probabilities in parentheses). Correlations are positive (+), negative (−) or not significant (NS). 1The relationship between sea urchin density and macroalgae is negative with all West-shallow survey sites but NS with West-shallow sea urchin survey sites only.

|  | Sea urchin density (ind. m<sup>-2</sup>) | % coral cover | % macroalgae cover | Parrotfish density (ind. 100 m<sup>-2</sup>) | % CCA cover | % turf algae cover |
| --- | --- | --- | --- | --- | --- | --- |
| Sea urchin density (ind. m<sup>-2</sup>) |  | **+** (Fig. 5a) | **-/NS<sup>1</sup>** (Fig. 5b) | **-** (Fig. 5c) | **NS** (p=0.3) | **NS** (p=0.4) |
| % coral cover |  |  | **-** (Fig. 6a) | **-** (Fig. 6b) | **+** (Fig. 6c) | **NS** (p=0.68) |
| % macroalgae cover |  |  |  | **NS** (p=0.06) | **-** (Fig. 6d) | **-** (Fig. 6e) |
| Parrotfish density (ind. 100 m<sup>-2</sup>) |  |  |  |  | **NS** (p=0.15) | **NS** (p=0.9) |
| % CCA cover |  |  |  |  |  | **NS** (p=0.25) |
| % turf algae cover |  |  |  |  |  |  |

The rest of the regression analyses not involving sea urchin density as a predictor *per se* were performed using all the survey data from all the sites. Among all sites, percent macroalgae cover was inversely related to percent coral cover (Fig. 6a). Parrotfish density was inversely related to percent coral cover (Fig. 6b, p=0.013), but not significantly related to percent macroalgae cover (Table 2, p=0.06), percent CCA cover (Table 2, p=0.15) and percent turf algae cover (Table 2, p=0.9). There was a positive correlation between percent coral cover and percent CCA cover (Fig. 6c, p=0.009) but no relationship between percent coral cover and percent turf algae cover (Table 2, p=0.684). Percent macroalgae cover was inversely related to both percent CCA cover (Fig. 6d, p=0.027) and percent turf algae cover (Fig. 6e, p<0.0001). Finally, there was no relationship between percent turf algae cover and percent CCA cover (Table 2, p=0.246). Note that percent turf algae cover was greater than percent CCA cover in all survey sites (Table 1). Table 2 summarizes the relationships among all the different variables.

**Figure 6.**
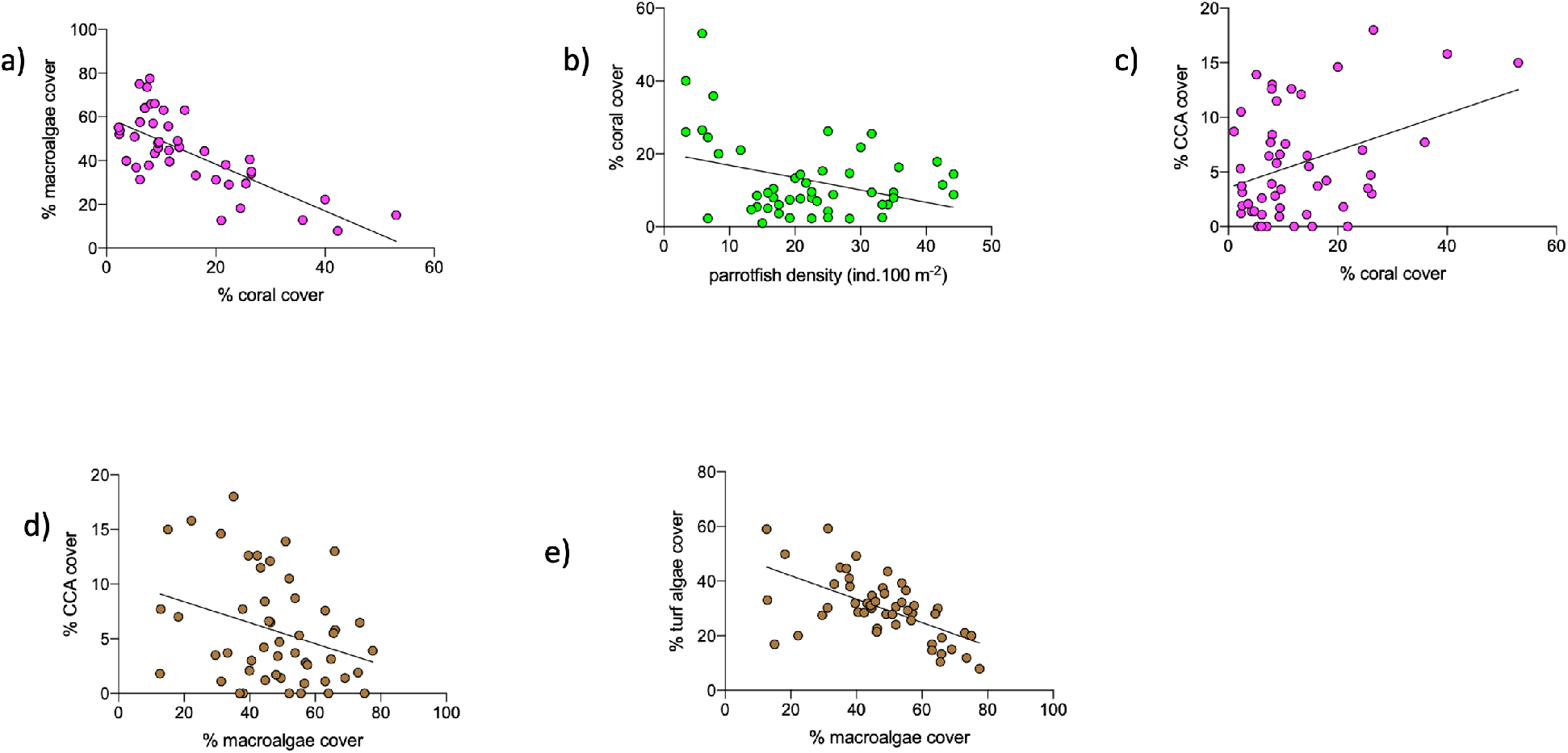
Significant relationships among different coral reef diversity measures in all 50 surveys from the 5 different site categories (West-shallow with urchins, West-shallow no urchins, West-deep no urchins, North-shallow no urchins, North-deep no urchins). a) Relationship between percent coral cover and percent macroalgae cover (p<0.0001). b) Relationship between parrotfish density and coral cover (p=0.013). c) Relationship between percent coral cover and percent CCA cover (p=0.009). d) Relationship between percent macroalgae cover and percent CCA cover (p=0.027). e) Relationship between percent macroalgae cover and percent turf algae cover (p<0.0001).

### Parrotfish size distribution in different survey sites

In all survey sites with the exception of the sea urchin survey sites, the most abundant parrotfish was the smallest size category (<11 cm) of *Sc. iseri*, the striped parrotfish (Fig. 7). In contrast, the most abundant parrotfish in the sea urchin survey sites was the larger size category (11-20 cm) of *Sc. taeniopterus*, the princess parrotfish. The lower overall parrotfish abundance of the sea urchin survey sites is also reflected in Figure 4c.

**Figure 7.**
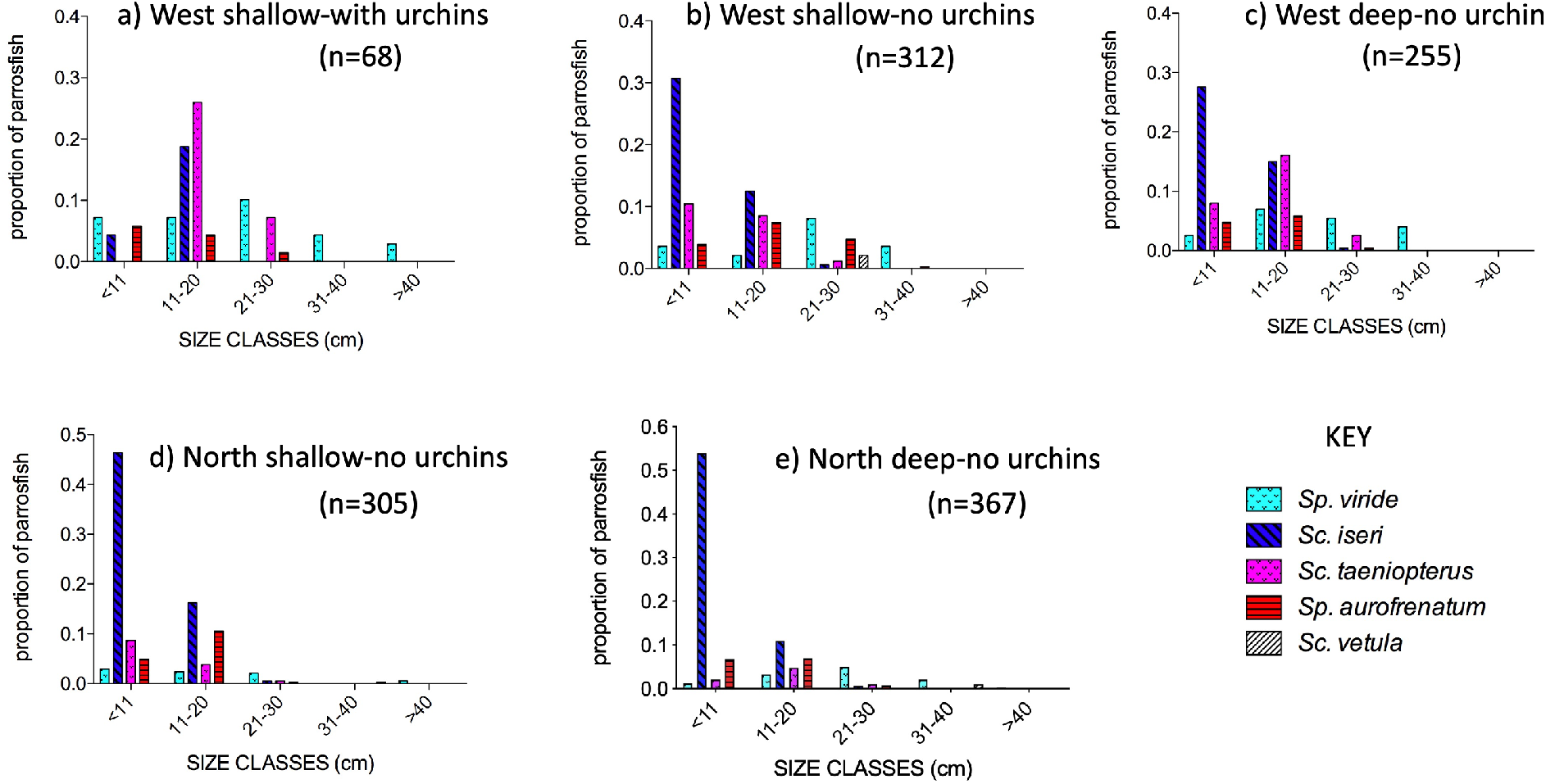
Proportion of parrotfish size categories and species distribution in the 5 different categories of survey sites. Numbers in parentheses are total fish count in the survey sites.

## DISCUSSION

### Significance of sea urchins

One of the main findings of this study is that the presence of sea urchins was associated with survey sites having higher levels of coral cover, lower levels of macroalgae cover, and lower densities of parrotfish than survey sites without sea urchins (Figs. 4a, b, c). Moreover, I found that coral cover and macroalgal cover were inversely related (Fig. 6a), a finding consistent with prior reports that coral and macroalgae compete with each other (Arnold, Steneck, and Mumby, 2010; Box and Mumby, 2007; Foster, Box, and Mumby, 2008; Jackson et al., 2014; Thurber et al., 2012). In fact, the data from Figure 6a could be superimposed on those of Figure S4 from Lester et al. (2020) regressing percent macroalgae cover on percent coral cover from 328 sites across the Caribbean from AGGRA surveys. Previous studies also have reported that areas with *Diadema* present were associated with areas of greater coral cover (Carpenter and Edmunds, 2006; Edmunds and Carpenter, 2001; Idjadi, Haring, and Precht, 2010), and like Bodmer et al (2015) my results revealed that sea urchin density was positively correlated with percent coral cover (Fig. 5a). The range of sea urchin density in this study was 0-0.75 individuals m^−2^, much lower than pre-die off densities of 20 or more individuals m^−2^ (Hughes et al., 2010). I was surprised, therefore, that such a small range of low densities could be related to such a large variation in coral cover of 2.2-53%. But both Williams et al. (2016) and Myhre and Acevedo-Guitierrez (2007) reported a similar positive relationship between coral cover and sea urchin density, at similar magnitudes. It is tempting to infer a causal relationship between the presence of *D. antillarum* and coral cover, the putative mechanism being grazing by sea urchins of the macroalgae that compete with corals as suggested by several studies (Carpenter and Edwards, 2006; Edmunds and Carpenter, 2001; Idjadi, Haring, and Precht, 2010). And survey sites with sea urchins in the present study were associated with lower levels of macroalgae cover (Fig. 4b). Indeed, experimental field studies in which *Diadema* were transplanted to various reef areas resulted in decreases in macroalgae following those transplants (Maciá, Robinson and Nalevanko, 2007; Williams, 2018). Maciá, Robinson and Nalevanko (2007) specifically noted a decrease in the macroalga *Dictyota*, the most abundant macroalga in their study sites in Jamaica and mine in Grand Cayman. A laboratory food preference study indicated that *Diadema* did ingest *Dictyota* (Solandt and Campbell, 2001). However, if sea urchins were facilitating greater coral cover due to their removal of macroalgae in Grand Cayman, I would have expected that sea urchin density would be inversely related to macroalgal cover as Bodmer et al (2015) reported. This was the case when all the West-shallow survey data were used (i.e. West-shallow survey sites with and without sea urchins) but when the regression analysis was repeated using data only from the sites with sea urchins, sea urchin density was not a predictor of macroalgal cover (Fig. 5b). Moreover, other studies found little association between macroalgal cover and the presence of *Diadema* (Lacey, Fourqurean and Collada-Vides, 2013; Mercado-Molina et al., 2015; Rodríguez-Barreras et al, 2018; Steiner and Williams, 2006). So while the lower percent macroalgae cover and higher percent coral cover are consistent with a causal relationship between those benthic measures and the presence of sea urchins, a causal connection cannot necessarily be confirmed by my results.

### Significance of parrotfish

Parrotfish density was inversely related to coral cover (Fig. 6b) and unrelated to macroalgal cover (Table 2), results in sharp contrast to several prior studies (Steneck, Arnold, and Mumby, 2014; Mumby, 2009; Mumby et al, 2006; Burkepile and Hay, 2010; Suchley and Alvarez-Filip, 2017). Those studies suggested that parrotfish grazed on macroalgae whose removal from the reef permitted the growth of corals. However, other reports found no relationship between parrotfish density and macroalgal cover (Francis et al, 2019; Russ, Questel, and Alcal, 2015; Suchley, McField, and Alvarez-Filip, 2016) and Clements et al. (2017) argued that the available evidence simply does not support parrotfish as primary consumers of macroalgae. In fact, Pattengill-Semmens and Semmens (2003) reported a positive relationship between parrotfish density and macroalgal cover in the Cayman Islands. Moreover, Onufrky et al (2018) reported a reciprocal relationship between *Diadema* density and the density of herbivorous fish, consistent with my finding that survey sites with sea urchins were associated with lower parrotfish densities than sites without sea urchins (Fig. 4c). Thus, the data presented here are not consistent with a causal relationship between parrotfish grazing, macroalgae removal, and concomitant growth of coral.

The species diversity and size distribution of parrotfish were different between the survey sites with and without sea urchins (Fig. 7). In survey sites without sea urchins, the smallest size category of striped parrotfish, *Sc. iseri*, was the most abundant, but in the sea urchin sites, a larger size category of princess parrotfish, *Sc. taeniopterus* was most abundant. Both of these species are small-bodied parrotfish that typically graze turf algae (Adam et al, 2015; Bonaldo, Hoey, and Bellwood, 2014), although my study did not reveal a relationship between turf algae cover and parrotfish density (Table 2). Fishing tends to disproportionately remove larger parrotfish, and thus smaller parrotfish tend to be less responsive to fishing pressure (Bonaldo, Hoey, and Bellwood, 2014). Shantz, Ladd and Burkpile (2020) reported that across the Caribbean, parrotfish assemblages are disproportionately represented by fish smaller than 11 cm, in agreement with the data in my study (Fig. 7) and Seemann et al (2018) noted that the parrotfish of greatest abundance by far in Caribbean Panama was *Sc. iseri*, again, in agreement with the data in my study. Although the removal of parrotfish due to fishing in Grand Cayman has been observed only recently (B. Johnson, Cayman Islands Department of Environment, pers. comm.) the prevalence of the small *Sc. iseri* and the low abundance of the larger bodied *S. viride* in my study are consistent with the removal of these larger parrotfish. Moreover, while the abundance of large (>31 cm) *S. viride* was low in all survey sites, the highest abundance was in the West-shallow sites with sea urchins (Fig. 7), having the lowest macroalgal cover (Fig. 4b), consistent with the report that macroalgal cover was inversely related to the density of large parrotfish across the Caribbean (Shantz, Ladd and Burkpile, 2020). However, none of my results derive from experimental manipulations so the finding that sites with large parrotfish are associated with lower macroalgal cover as are sites with sea urchins (Fig. 4b), while consistent with a putative mechanism of algal removal by these nominal herbivores, cannot be taken as confirmation. Moreover, my data do not permit distinguishing whether the presence of large parrotfish or the presence of sea urchins or both are responsible for the lower macroalgal cover. In fact, Tebbett et al, (2020) argued that the presence and abundance of a particular herbivore species is not a predictor of its functional role in Caribbean coral reef ecosystems. They demonstrated that in spite of their abundance, parrotfish contributed very little to the ecosystem function of macroalgal removal.

### Can different studies be reconciled?

The measures of reef biodiversity reported here (Table 1) agree with those reported in prior studies. For example, the coral cover of roughly 31% in survey sites with sea urchins and roughly 10% in sites without sea urchins I reported agree with a recent report of 17% coral cover on Grand Cayman (Manfrino and Dell, 2019). Similarly, that study reported macroalgal cover of 57% in Grand Cayman compared to the roughly 46-58% macroalgal cover that I recorded in sites without sea urchins (Table 1). The density of sea urchins I found in Grand Cayman of 0-0.75 individuals m^−2^ with an average density of 0.32 individuals m^−2^ in the survey sites with urchins is consistent with reports from all over the Caribbean, summarized by Lessios (2016), of average densities of 0.02-1.43 individuals m^−2^. Parrotfish densities (in survey sites without sea urchins) of 20-30 individuals 100 m^−2^ are similar to those reported by Manfrino and Dell (2019) and the size distributions of parrotfish I reported here are similar to those reported in other studies (Bruckner, 2010; Adam et al, 2018; Shantz, Ladd and Burkepile, 2020). The agreement in the *magnitude* of reef biodiversity measures should dispel concern, to some extent, that the varying methods used in the different studies might have introduced important variables, but it also begs the question as to why the *relationships* among the various biodiversity measures differ so much across different studies. Why, for example, do so many studies report a positive, potentially causal, relationship between parrotfish abundance and coral cover (Burkepile and Hay, 2010; Shantz, Ladd, and Burkepile, 2020; Steneck, Arnold, and Mumby, 2014), while other studies, including mine, show no such relationship (Suchley, McField, and Alvarez-Filip, 2016; Tebbett et al, 2020) or even dispute the claim of parrotfish macroalgae grazing (Clements et al., 2017)? In fact, in my study, parrotfish abundance was inversely related to coral cover, suggesting, if anything, corallivory (Fig. 6b). Different studies have suggested that the category of “parrotfish” is too broad to be a useful explanatory variable because there is so much diversity of size and behavior both within and across different species (Ruttenberg et al., 2019; Steneck, Arnold, and Mumby, 2014). In fact, Ruttenberg et al (2019) and Lefcheck et al (2019) suggested that there is little redundancy among herbivores at the scale of individual reefs for the function of macroalgae removal and turf algae removal, respectively. They argued that studies that characterize herbivory at a regional scale do not provide sufficient resolution to assess the variation in herbivory in individual reefs within a region. My results are consistent with that finding in that the coral reef survey sites in which I found sea urchins might well have been overlooked simply because they were in such a small area. Bodmer et al (2015) also reported a sequestered population of *Diadema* at unusually high densities in a newly discovered reef system off Honduras. This reef system was geographically separated from the nearest Caribbean reef system of Utila by roughly 60 km, the Utila reefs housing the typically low sea urchin densities of the rest of the Caribbean. But the Western-shallow reef area in Grand Cayman that exhibited the higher *Diadema* densities in my study was contiguous with survey areas without sea urchins. A diver could easily see and swim from reefs with *Diadema* to reefs without, within 15-30 m of each other. There were no apparent barriers that would interfere with broadcast gamete movement of *Diadema*. However, the shortest distance between reefs with and without sea urchins was greater than the 3 m beyond which Levitan (1991) reported that gamete dilution made the probability of fertilization unlikely. So the scale at which there was significant variation in biodiversity measures in Grand Cayman was on the order of 15-30 m.

Most studies on *Diadema* density in the Caribbean have noted the absence of recovery of these populations to pre-die off levels (Bodmer et al., 2015; Lessios, 2016; Tuohy, Wade, and Weil, 2020). In reefs where some recovery has been noted, however, the greater relative habitat complexity of those reefs might provide shelter to juvenile sea urchins facilitating their growth and persistence (Bodmer et al, 2015; Maciá, Robinson and Nalevanko, 2007; Tuohy, Wade, and Weil, 2020). I did not directly measure the rugosity of the reefs in this study, but I did note that the survey sites with sea urchins were typically smaller patches of discrete reef mounds compared to the longer reef fingers covered with *Dictyota* nearby found at the same depths. This does not explain, however, the absence of sea urchins in other such discrete patches at similar depths in this study. Nevertheless, those sites could be colonized in the future. Perhaps sea urchins were grazing on macroalgae on those discrete mounds or perhaps, for some other reason, the macroalgae levels were low on the discrete reef mounds, which attracted sea urchins, or at the very least, permitted them to persist due to the habitat complexity.

Both CCA cover and turf algae cover were inversely related to macroalgae cover (Fig. 16a, b) consistent with prior reports (Adam et al, 2018; Tebbett and Bellwood, 2019). Moreover, CCA cover and coral cover were positively related (Fig. 6c), again, consistent with the observation that CCA plays an important role in coral settlement and persistence (Tebbett and Bellwood, 2019). I found no evidence that CCA and turf algae actively compete with each other, however, as their benthic covers were unrelated (Table 2).

### Marine Protected Areas

The MPA in Grand Cayman was established on the western side of the island in 1986 (Fig. 1). McCoy, Dromard, and Turner (2010) compared fish communities within the MPA to non-MPA communities more than 20 years after the establishment of the MPA. They reported that fish biomass of various functional groups, including herbivores was greater inside the MPA. The density of the herbivores, however, did not differ according to location, consistent with my findings, more than 30 years after the establishment of the MPA in Grand Cayman. They also found that there was a greater proportion of larger parrotfish within the MPA, whereas in my study, the only difference in size distribution of parrotfish was between the sea urchin survey sites (within the MPA) and all the other sites, some of which were within the MPA (Fig. 7).

Just as with the varying relationships among biodiversity metrics of reefs, there is no consistent evidence across studies as to the efficacy of MPAs in promoting coral reef protection (Suchley and Alvarez-Filip, 2018; Woodcock et al., 2016). One of the difficulties in comparing different MPAs is that there is no consistent description, implementation, and enforcement of MPAs (Roberts et al., 2017; Strain et al., 2019). For example, some MPAs are no-take zones while others, like Grand Cayman permit some extractive fishing practices. And while fish diversity has been used as an indicator of reef health, the most significant measure of reef persistence and resilience is scleractinian coral cover. Larger and older MPAs appear to be the most effective in coral protection (Strain et al., 2019). I found no difference in coral cover between sites (without sea urchins) within the Grand Cayman MPA compared to non-MPA sites (Fig. 4a). Many researchers have argued that management of herbivorous fish populations within MPAs is insufficient to oppose the overwhelming assaults of climate change and growing human populations in coastal areas (Bruno, Côté, and Toth, 2019; Graham, et al., 2020).

## Conclusions

Grand Cayman is not an outlier of Caribbean coral reefs in terms of measures of coral reef diversity and relationships among diversity variables. But there is so much heterogeneity among those variables across Caribbean reefs that no one reef or even region can be considered an exemplar. Some studies, like mine, have reported associations between coral cover and *Diadema* density, while others have reported relationships between parrotfish abundance and coral cover, and still others have found that neither sea urchins nor parrotfish were predictors of coral cover. This heterogeneity is likely due to the responses of reefs to particular local stressors and global stressors and their interactions (Benkwitt, Wilson, and Graham, 2020; Bruno, Côté, and Toth, 2019; Chollett et al., 2017). This combination of site-specific conditions and responses to stressors makes the search for a single variable or two, e.g. herbivorous fish or invertebrates, to help scientists and managers predict the trajectory and resilience of reefs unlikely.

Grand Cayman is typical of Caribbean islands that have experienced and will continue to experience increases in coastal development and human population pressures, both enormous threats to coral reefs (Friedman et al., 2020; Reigl and Glynn, 2020). The MPAs that appear to offer some protection against reef degradation are characterized by strict enforcement of restrictions on extraction of reef resources (Roberts et al., 2017). Nevertheless, these efforts are not likely to permit the persistence and resilience of reefs as we know them now due to climate change (Benkwitt, Wilson, and Graham, 2020; Graham et al., 2020; Suchley and Alvarez-Filip, 2018). Present day reefs are already substantially degraded compared to 40 years ago (Hughes et al, 2017). The baselines of reef diversity keep changing, with decreases in hard coral cover and increases in macroalgal cover. Reefs are becoming less structurally complex as dead carbonate skeletons are worn away, with concomitant changes in the composition of organisms that live there. So what does resilience mean? As more stressors impact reefs and corals die off, do we want to “manage” reefs for a return to the already depauperate composition of less than 10% coral cover? Bruno, Côté, and Toth (2019) reported that there was little evidence to support the notion of “managed resilience.” I concur with myriad prior reports and virtually all coral reef scientists of whom I am aware, that if we do nothing to rein in coastal development and climate change, reefs even as we know them now, in their degraded states, will perish (e.g. Bruno, Côté, and Toth, 2019; Graham et al., 2020; Hughes et al, 2017; Suchley and Alvarez-Filip, 2018).

## Supporting information

Supplementary material Sherman

chain no urchins video

chain with urchins video

## Acknowledgements

Thanks to Tim Austin, Deputy Director of the Cayman Islands Department of Environment, for ongoing advice, encouragement, and assistance. Additionally, thanks go to John Bothwell and Director Gina Ebanks-Petrie of the Cayman Islands Department of Environment. Dusty Norman and his staff at DNS Diving provided valuable technical assistance for which I am grateful. Thanks to Katie Alpers of Indigo Divers, sea urchin finder extraordinaire. Kerry Woods and Kathryn Montovan provided helpful comments on the manuscript. Finally, thanks to my underwater research assistants Arlene Wyman, Elizabeth Schappel, and Hannah Leigh.

