## Supplementary material Sherman for "Sea urchins, parrotfish and coral reefs in Grand Cayman, BWI: exemplar or outlier?"

Elizabeth Sherman

Supplementary Material

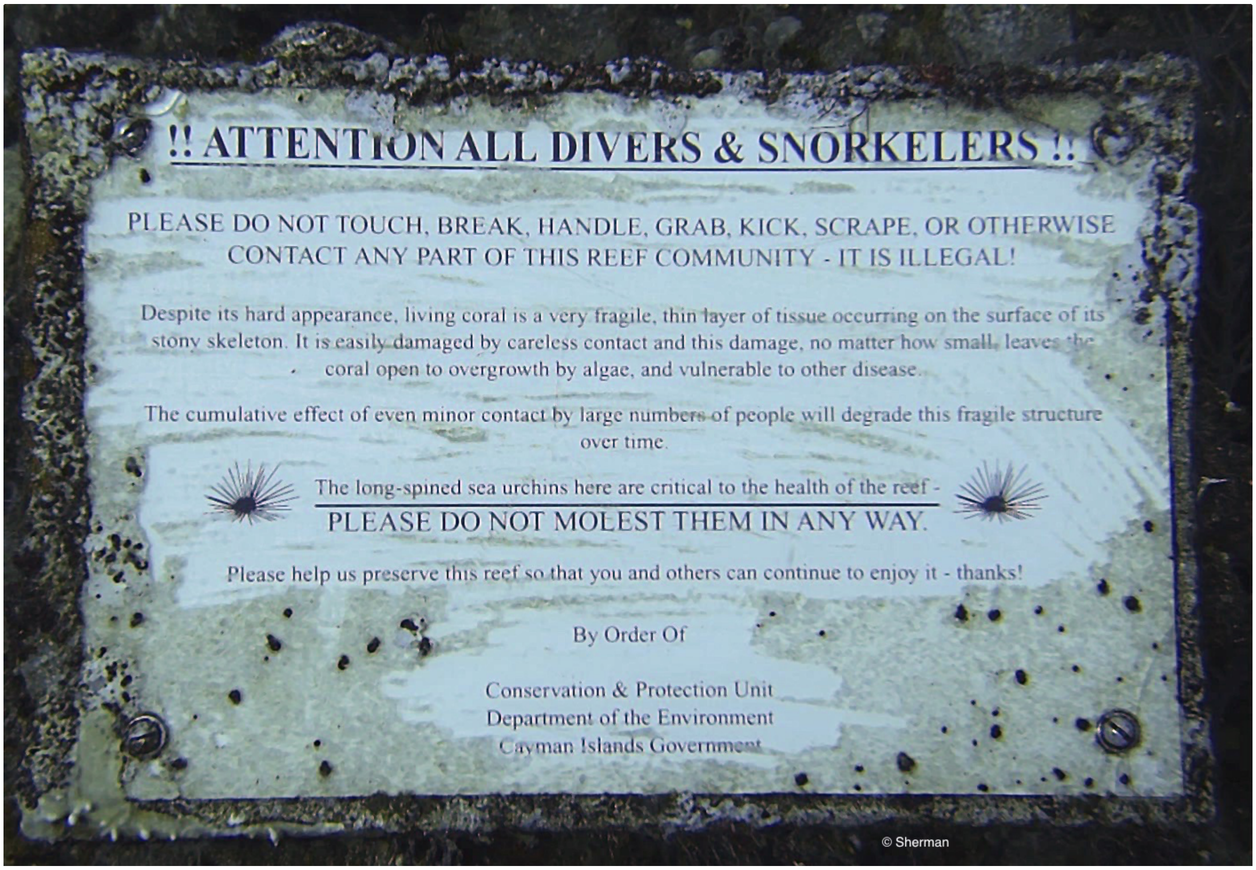


**Figure 1S.** Photo of a sign at a depth of roughly 18 m at the pin of a mooring site in Grand Cayman, cautioning divers not to disturb the sea urchin. The sign was likely placed by the Department of Environment prior to 1990 (pers. comm. Tim Austin, Cayman Islands Department of Environment) suggesting that at one time, there had been observable sea urchin populations at that depth.

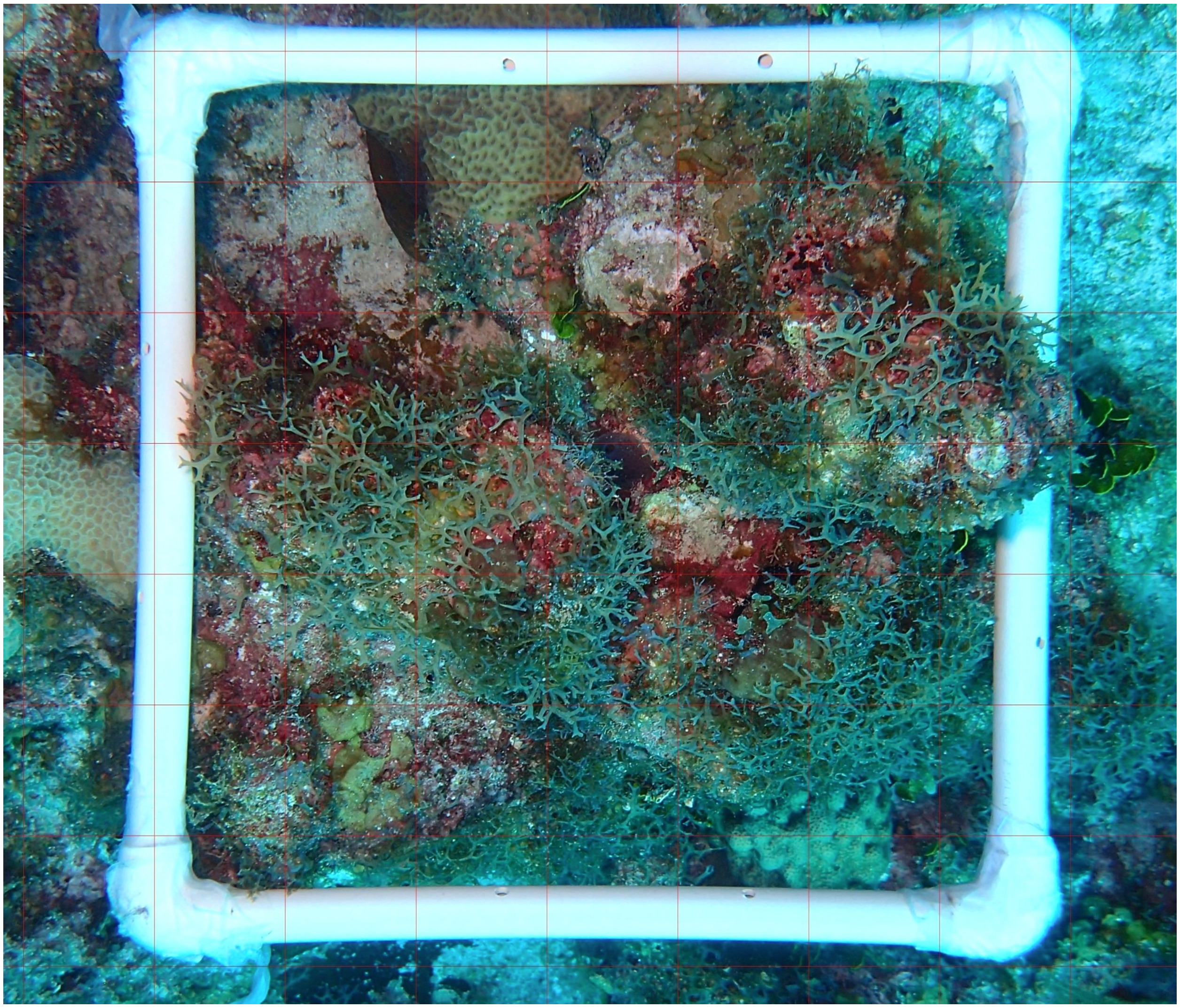

**Figure 2S.** Example of transect photo with superimposed 6 x 6 grid (faint lines in red).

**Two-way ANOVAS**

Initial 2-way ANOVAs to determine if data from survey sites with sea urchins (West-shallow with urchins) could be treated as a separate site category along with the other 4 site categories: West-shallow no urchins, West-deep no urchins, North-shallow no urchins, North deep no urchins.

<u>Coral Cover</u>

Table 1Sa. When the survey sites with sea urchins were excluded from the two-way ANOVA, there was no significant difference in percent coral cover among all the survey sites, and neither year nor interaction was significant.

| ANOVA TABLE | df | F | p |
|---|---|---|---|
| **Interaction** | 6 | 0.772 | 0.5972 |
| **Year** | 2 | 0.062 | 0.940 |
| **Survey site** | 3 | 1.24 | 0.311 |
| **Residual** | 32 | | |

Table 1Sb. When the sea urchin survey site data were added to the West-shallow survey site data and the two-way ANOVA was repeated, the survey site category became significant, while year and interaction remained not significant. Thus it was legitimate to perform a subsequent one-way ANOVA (with pooled years) using the percent coral cover from the sea urchin survey sites as a separate site category (West-shallow with urchins, see Fig. 4).

| ANOVA TABLE | df | F | p |
|---|---|---|---|
| **Interaction** | 6 | 0.109 | 0.995 |
| **Year** | 2 | 0.029 | 0.971 |
| **Survey site** | 3 | 2.91 | 0.046 |
| **Residual** | 41 | | |

Macroalgal cover

Table 2Sa. When the survey sites with sea urchins were excluded from the two-way ANOVA, there was no significant difference in percent macroalgal cover among all the survey sites, and neither year nor interaction was significant.

| ANOVA TABLE | df | F | p |
|---|---|---|---|
| **Interaction** | 6 | 1.37 | 0.279 |
| **Year** | 2 | 2.34 | 0.135 |
| **Survey site** | 3 | 2.29 | 0.279 |
| **Residual** | 30 | | |

Table 2Sb. When the sea urchin survey site data were added to the West-shallow survey site data and the two-way ANOVA was repeated, the survey site category became significant, while year and interaction remained not significant. Thus percent macroalgae cover from the sea urchin survey sites as a separate site category (West-shallow with urchins, see Fig. 5).

| ANOVA TABLE | df | F | p |
|---|---|---|---|
| **Interaction** | 6 | 0.662 | 0.681 |
| **Year** | 2 | 2.04 | 0.10 |
| **Survey site** | 3 | 6.10 | 0.0016 |
| **Residual** | 41 | | |

<u>Parrotfish density</u>

Table 3Sa. While parrotfish density is potentially a predictor of coral reef cover and macroalgal cover, I performed a two-way ANOVA testing whether site or year affected parrotfish density. When the survey sites with sea urchins were excluded from the two-way ANOVA, there was no significant difference in parrotfish density among all the survey sites, and neither year nor interaction was significant.

| ANOVA TABLE | df | F | p |
|---|---|---|---|
| **Interaction** | 6 | 0.662 | 0.636 |
| **Year** | 2 | 2.85 | 0.10 |
| **Survey site** | 3 | 0.997 | 0.41 |
| **Residual** | 31 | | |

Table 3Sb. When the sea urchin survey site data were added to the West-shallow survey site data and the two-way ANOVA was repeated, the survey site category became significant, while year and interaction remained not significant. Thus it was legitimate to perform a subsequent one-way ANOVA (with pooled years) using parrotfish density from the sea urchin survey sites as a separate site category (West-shallow with urchins, see Fig. 6).

| ANOVA TABLE | df | F | p |
|---|---|---|---|
| **Interaction** | 6 | 0.36 | 0.899 |
| **Year** | 2 | 2.073 | 0.14 |
| **Survey site** | 3 | 2.803 | 0.05 |
| **Residual** | 40 | | |

CCA cover

Table 4Sa. When the survey sites with sea urchins were excluded from the two-way ANOVA, there was no significant difference in percent CCA cover among all the survey sites, and neither year nor interaction was significant.

| ANOVA TABLE | df | F | p |
|---|---|---|---|
| **Interaction** | 6 | 2.07 | 0.087 |
| **Year** | 2 | 1.490 | 0.242 |
| **Survey site** | 3 | 1.486 | 0.238 |
| **Residual** | 30 | | |

Table 4Sb. When the sea urchin survey site data were added to the West-shallow survey site data and the two-way ANOVA was repeated, the survey site category became significant, while year remained not significant. However, the interaction between survey site and year was significant. And with that caveat, I did perform a subsequent one-way ANOVA.

| ANOVA TABLE | df | F | p |
|---|---|---|---|
| **Interaction** | 6 | 3.637 | 0.006 |
| **Year** | 2 | 1.279 | 0.2899 |
| **Survey site** | 3 | 3.785 | 0.0181 |
| **Residual** | 38 | | |

Turf algae cover

Table 5Sa. When the survey sites with sea urchins were excluded from the two-way ANOVA, there was no significant difference in percent turf algae cover among all the survey sites, and while survey site was not significant, year was significant, making the legitimacy of pooling data from the different years problematic.

| ANOVA TABLE | df | F | p |
|---|---|---|---|
| **Interaction** | 6 | 0.5887 | 0.7366 |
| **Year** | 2 | 7.436 | 0.0024 |
| **Survey site** | 3 | 1.735 | 0.1809 |
| **Residual** | 30 | | |

Table 5Sb. When the sea urchin survey site data were added to the West-shallow survey site data and the two-way ANOVA was repeated, the survey site category remained not significant, while year remained significant, so I did not perform a subsequent one-way ANOVA.

| **ANOVA TABLE** | **df** | **F** | **p** |
|---|---|---|---|
| **Interaction** | 6 | 0.5899 | 0.7362 |
| **Year** | 2 | 5.509 | 0.0079 |
| **Survey site** | 3 | 2.360 | 0.0867 |
| **Residual** | 38 | | |
